# Differential roles of pyramidal and fast-spiking, GABAergic neurons in the control of glioma cell proliferation

**DOI:** 10.1101/2020.03.05.978825

**Authors:** Elena Tantillo, Eleonora Vannini, Chiara Cerri, Cristina Spalletti, Antonella Colistra, Chiara Maria Mazzanti, Mario Costa, Matteo Caleo

## Abstract

Recent studies have demonstrated an active role for neurons in glioma progression. Specifically, peritumoral neurons establish functional excitatory synapses with glioma cells, and optogenetic stimulation of cortical pyramidal neurons drives tumor progression. However, the specific role of different subsets of cortical neurons, such as GABAergic interneurons, remains unexplored. Here, we directly compared the effects of optogenetic stimulation of pyramidal cells vs. fast-spiking, GABAergic neurons. In mice inoculated with GL261 cells into the motor cortex, we show that optogenetic stimulation of pyramidal neurons enhances glioma cell proliferation. In contrast, optogenetic stimulation of fast-spiking, parvalbumin-positive interneurons reduces proliferation as measured by BrdU incorporation and Ki67 immunolabelling. Since both principal cells and fast-spiking interneurons are directly activated by sensory afferent input, we next placed tumors in the occipital cortex to test the impact of visual stimulation/deprivation. We report that total lack of visual input via dark rearing enhances the density of proliferating glioma cells, while daily visual stimulation by gratings of different spatial frequencies and contrast reduces tumor growth. The effects of sensory input are region-specific, as visual deprivation has no significant effect on tumor proliferation in mice with gliomas in the motor cortex. We also report that sensory stimulation combined with temozolomide administration delays the loss of visual responses in peritumoral neurons. Altogether, these data demonstrate complex effects of different neuronal subtypes in the control of glioma proliferation.

**Highlights:**

- Activity of GABAergic neurons reduces glioma cell proliferation
- Levels of sensory afferent input regulate tumor proliferation
- Effects of sensory input are region-specific

## INTRODUCTION

Gliomas are malignant primary brain tumors with a poor outcome. Glioblastoma multiforme (GBM) is the most aggressive form, belonging to the class IV of the World Health Organization classification(Burnet et al., 2007; Perry et al., 2016). Despite advances in the last years(Ozdemir-Kaynak et al., 2018; Reitman et al., 2018), GBM remains one of the most difficult cancer to treat, with a median survival of 15 to 17 months and 5-year overall survival of 5%(Ostrom et al., 2016). Moreover, all the approved experimental treatments for high-grade glioma patients do not offer long-term benefits in symptom improvement or quality of life(Laub et al., 2018; Wick et al., 2018). This devastating scenario strongly indicates a lack of knowledge in the biology of glioma and in its interaction with the peritumoral areas. Therefore, the development of novel therapies requires a better understanding of the communication between the tumor and resident brain cells.

Evidences of highly disabling peritumoral dysfunctions in patients with glioma(van Kessel et al., 2017) and experimental data from literature(Armstrong et al., 2016; Buckingham and Robel, 2013) suggest that glioma cells exert a powerful influence on neighbouring neurons. For example, it has been demonstrated that glioma cells extrude high amounts of glutamate resulting in excitotoxicity and tumor invasion(Marcus et al., 2010; Rzeski et al., 2001; Sontheimer, 2008). Infiltrating glioma cells also perturb chloride homeostasis in pyramidal neurons, with dowregulation of KCC2 (K-Cl cotransporter 2) expression and consequent excitatory actions of gamma-aminobutyric acid (GABA)(Campbell et al., 2015; Conti et al., 2011; Pallud et al., 2014). Glioma cells harbouring specific genetic mutations drive synaptic changes in the surrounding neuronal microenvironment, specifically an enhancement in the density of excitatory boutons coupled with a decrease in inhibitory puncta (Yu et al., 2020). Altogether, these alterations render peritumoral neuronal networks hyper-excitable and more prone to seizures (Buckingham et al., 2011; Venkatesh et al., 2019).

On the other hand, there is now robust evidence that neuronal activity affects glioma cell behaviour. The most relevant studies of the latest years have demonstrated that pyramidal cell activity promotes glioma proliferation through secreted factors, in particular neuroligin-3 (NLGN3) (Venkatesh et al., 2015, 2017). Indeed, optogenetic stimulation of excitatory, pyramidal neurons enhances glioma proliferation via the release of NLGN3 into the tumour microenvironment (Venkatesh et al., 2015, 2017). Moreover, it has been recently discovered that peritumoral neurons and glioma cells directly interact through AMPA-mediated synapses that drive tumor growth and invasion (Venkataramani et al., 2019; Venkatesh et al., 2019).

Cortical circuitry consists of several subtypes of neurons, varying in their molecular signatures, electrophysiological properties, and connectivity patterns. In particular, about 20% of cortical neurons are GABAergic inhibitory cells, which are further subdivided into at least three populations: (i) fast-spiking, parvalbumin-positive cells which synapse on soma and axon hillock of pyramidal neurons; (ii) somatostatin-positive interneurons characterized by a preferential dendritic targeting of their synapses; (iii) interneurons that express the serotonin receptor 5HT3aR, comprising those releasing vasoactive intestinal peptide (VIP) (Lim et al., 2018).

Whether and how the activity of these interneurons modulates glioma proliferation is currently unknown. To begin addressing this issue, here we have tested the impact of a selective optogenetic stimulation of fast-spiking, parvalbumin-positive interneurons. These cells are particularly important in the control of cortical excitability due to their perisomatic synapses which potently regulate the firing properties of principal neurons (Deidda et al., 2015; Di Cristo et al., 2004).

Pyramidal cells and fast-spiking interneurons in the neocortex respond to afferent sensory stimulation, albeit with different dynamics and selectivities (Hofer et al., 2011; Runyan and Sur, 2013). Fast-spiking interneurons play a key role in sensory processing as they are directly contacted by thalamocortical afferents (Sugiyama et al., 2008; Yu et al., 2016). It is therefore of interest to determine whether sensory input impacts on glioma proliferation, potentially via its effects on the activity of principal cells and interneurons.

Here, we use the widely employed GL261 syngeneic mouse model of high grade glioma(Miyai et al., 2017; Szatmári et al., 2006; Vannini et al., 2016, 2017) to directly compare the effects of optogenetic stimulation of pyramidal cells vs. fast-spiking, GABAergic interneurons. We show that the activity of these neuronal populations has opposite effects on the proliferation rate of adjacent glioma cells. We also test the impact of visual afferent input on the proliferation of tumors placed in the occipital cortex.

## MATERIALS AND METHODS

### Animals

Adult (age comprised between postnatal day 60 and 120) mice were used for this study. We employed (i) wild type C57BL/6J, (ii) Thy1-ChR2 (B6.Cg-Tg (Thy1-ChR2/EYFP) 18Cfng/J, Jackson Laboratories, USA) and (iii) PV-Cre (Tanahira et al., 2009) (B6;129P2-Pvalb tm1(cre)Arbr/J, Jackson Laboratories, USA) mice in different experiments. Animals were bred in our animal facility and housed in a 12 hours light/dark cycle, with food and water available *ad libitum*. All experimental procedures comply with the ARRIVE guidelines. They were performed in conformity to the EU Council Directive 2010/63/EU and were approved by the Italian Ministry of Health.

### Tumor induction

Under avertin anesthesia (intraperitoneal injection of 2,2,2-tribromoethanol solution; 250 mg/kg body weight; Sigma Aldrich, USA), mice received a stereotaxically guided injection of 40,000 GL261 cells as described(Vannini et al., 2016, 2017). In C57BL/6J, Thy1-ChR2 and PV-Cre mice, cells were injected either into the primary visual cortex(Vannini et al., 2016) or into the motor cortex(Vannini et al., 2017) in correspondence with the forelimb representation (caudal forelimb area(Alia et al., 2016; Allegra Mascaro et al., 2019)).

### Optogenetic stimulation in Thy1-ChR2 mice

Thy1-ChR2 mice express the gene for channelrhodopsin-2 (ChR2) under the Thy1 promoter, mainly in deep cortical projection neurons (Arenkiel et al., 2007; Spalletti et al., 2017). In these mice, a portion of cranial bone around the point of injection was removed with a drill. After glioma injection into the motor cortex, a chamber of dental cement (Tetric EvoFlow, Ivoclar Vivadent, Switzerland) was created around the craniotomy and filled with a layer of agar (Sigma Aldrich, USA) and the silicone elastomer Kwik (World Precision Instrument, USA) to preserve the cortical surface. A metal post was placed on the occipital bone and fixed with dentistry cement (KERR, Dental Leader, Italy). Awake animals were subjected to a period of habituation to the apparatus before the stimulation session, made in awake mice.

After 13 days from tumor injection, a group of Thy1-ChR2 glioma-bearing mice was stimulated with an optogenetic fiber according to the single stimulation protocol described in Venkatesh et al., 2015 (Venkatesh et al., 2015). The tip of the optic fiber was positioned stereotaxically over the dura mater surface. Animals were stimulated with cycles of 473 nm light pulses at 20 Hz for 30 s, followed by 90 s of recovery over a 30 min period. Optogenetic stimulation was done using the setup described in Spalletti et al., 2017 (Spalletti et al., 2017). The control group was let in a head-fixed position without fiber stimulation in order to reproduce the same stress conditions of stimulated animals. Immediately before the beginning of the stimulation protocol, BrdU (5-Bromo-2′-deoxyuridine; Sigma Aldrich, USA) was intraperitoneally (i.p) administered (50 mg/kg) and animals were sacrificed for immunostaining after 24 hours.

### Adeno-associated virus (AAV) injection for ChR2 expression in Parvalbumin interneurons

One week before the GL261 graft, a cohort of PV-Cre and C57BL/6J mice as controls, received two stereotaxically guided injections of 600 nl of an AAV vector (AAV1.EF1.dflox.hChR2(H134R)-mCherry.WPRE.hGH (Addgene, USA)), one 500 μm anterior and one 500 μm posterior to the site of tumor injection into the motor cortex. PV-Cre mice express Cre recombinase in parvalbumin-expressing neurons. The AAV vector contains the channelrhodopsin-2 (ChR2) gene which is expressed in parvalbumin interneurons through Cre-mediated recombination. We refer to PV-ChR2 to indicate PV-Cre mice injected with the AAV. The two sites of injection were chosen in order to allow the expression of ChR2 in Parvalbumin interneurons located in the motor cortex surrounding the tumor mass. The surgery was performed under avertin anesthesia (intraperitoneal injection of 2,2,2-tribromoethanol solution; 250 mg/kg body weight; Sigma Aldrich, USA).

### Optogenetic stimulation of Parvalbumin interneurons

After the GL261 graft in PV-ChR2 mice, a portion of cranial bone anterior and posterior to the site of tumor injection was thinned with a drill and a chamber of dental cement (Tetric EvoFlow, Ivoclar Vivadent, Switzerland) was created around the craniotomy and filled with a layer of agar (Sigma Aldrich, USA) and the silicone elastomer Kwik (World Precision Instrument, USA) to preserve the cortical surface. A metal post was placed on the occipital bone and fixed with dentistry cement (KERR, Dental Leader, Italy) as previously described. Awake animals were subjected to a period of habituation to the apparatus before the stimulation sessions.

Starting 7 days from tumor injection, a group of PV-ChR2 mice was daily stimulated for 7 days with an optogenetic fiber positioned stereotaxically over the dura mater surface. Cycles of 15 min of 473 nm light pulses at 40 Hz (Cardin et al., 2009) for 3 s, followed by 90 s of recovery, were used to stimulate the anterior and posterior part of the craniotomy over a total period of 30 min. Optogenetic stimulation was done using the setup described in Spalletti et al., 2017 (Spalletti et al., 2017). The control group was let in a head-fixed position without fiber stimulation in order to reproduce the same stress conditions of stimulated animals. The day before the end of the protocol, BrdU (5-Bromo-2′-deoxyuridine; Sigma Aldrich, USA) was intraperitoneally administered (50 mg/kg) and animals were sacrificed for immunostaining after 24 hours.

### Manipulation of visual afferent input

C57BL/6J mice bearing tumors in the occipital cortex were subjected to different rearing conditions in order to manipulate afferent visual input from day 11 to day 14 post glioma injection. A group of animals was visually deprived through dark rearing (DR) in ventilated, completely light-tight shells, while another cohort of mice were stimulated with square wave and sinusoidal gratings (visual stimulation, VS) for 8 hours/day; during the stimulation, animals were put in a plexiglass cage surrounded by 4 LCD screens (one per side) with water and food available *ad libitum* and exposed to the visual stimulation protocol described below. Animals with glioma into the primary visual cortex that were maintained into the normal light/dark cycle were called standard light (SL). To test the region specificity of the effects of DR, a group of C57BL/6J mice bearing tumors in the motor cortex was reared in complete darkness as described above from day 11 to day 14 post glioma graft. BrdU (5-Bromo-2′-deoxyuridine; Sigma Aldrich, USA) was administered i.p. (50 mg/kg) on day 13 and animals were sacrificed after 24h for immunohistochemical analysis.

### Visual stimulation protocol

Visual stimuli were generated by a VSG2:5 card (Cambridge Research System, Cheshire, UK) and presented on the face of all the 4 LCD screens. The 8 hour protocol was composed by the repetition of two main modules of sinusoidal gratings of different spatial frequencies and contrasts: (i) a block of steady-state stimuli, consisting of gratings of increasing contrasts sinusoidally modulated at 2-6 Hz and (ii) a block of transient stimuli, i.e. gratings of increasing contrasts abruptly reversed at 1 Hz with a spatial frequency from 0.4 to 0.1 c/deg. Each block contained both horizontal and vertical gratings. The screen luminance was maintained at a low level (2 cd/m^2^) to minimize animals’ stress.

### Immunohistochemical analysis

Animals were deeply anesthetized with chloral hydrate and perfused transcardially with Phosphate Buffer Saline (PBS; Sigma Aldrich, USA) followed by fixative (4% paraformaldehyde, 0.1 M sodium phosphate, pH 7.4). Brains were gently removed, post-fixed for 4 h in the same fixative at 4°C, cryoprotected by immersion in 30% sucrose and cut using a sliding microtome (Leica, Germany) to obtain coronal sections of 50 μm of thickness.

To detect neuronal components infiltrating the tumor mass, brain sections were stained for the axonal marker Tau (1:1000; SySy, Germany) and for the neural marker NeuN (1:1000, Millipore, Germany). Thy1-ChR2 glioma-bearing mice were also stained for the neural marker Parvalbumin (1:500; SySy, Germany). For the evaluation of glioma cell proliferation, brain serial sections were stained with two markers of proliferation, Ki67 (1:400; Abcam, UK) and BrdU (1:500; Abcam, UK). Slices were incubated with fluorophore-conjugated secondary antibodies (Jackson Immunoresearch, USA) and with Hoechst dye (1:500; Sigma Aldrich, USA) for nuclei visualization.

In PV-ChR2 mice, we assessed colocalization of mCherry-ChR2 and parvalbumin. While mCherry staining was detectable in coronal sections without any amplification step, parvalbumin immunolabelling was performed with a primary guinea pig polyclonal antibody (1:500; SySy, Germany) as described (Busti et al., 2020). Counts of double labelled cells were performed using Neurolucida software (Allegra et al., 2017).

### Image acquisition and analysis of tumor cell proliferation

Fluorescent images were acquired using a Zeiss Axio Observer microscope equipped with Zeiss AxioCam MRm camera (Carl Zeiss MicroImaging GmbH, Germany). Slides were coded to ensure blinding during the acquisitions. Images were acquired through a 10x objective and joined precisely using Zen Blue Edition software (Carl Zeiss MicroImaging GmbH, Germany). High magnifications of neural components were acquired also through a 20x objective. For the quantification of the density of Ki67- or BrdU-positive cells, we acquired the whole area of the tumor for each coronal section. Images were processed using Image J software (National Institute of Health, USA). Density of proliferating cells was expressed as the fraction of area occupied by Ki67- or BrdU-positive cells with respect to total tumor area(Garofalo et al., 2015). For comparison between different treatments, at least 3 animals per group were analysed.

To minimize the variation between different immunostaining reactions and to normalize the data, we always processed a control group (i.e., reared in standard laboratory conditions and without any optogenetic stimulation) together with the experimental animals. Measurements of tumor proliferation were normalized to the mean of the control group in the same immunohistochemical reaction. Statistical comparison were performed by cumulating the data obtained from all sections in the same experimental group. For all experiments, we also report the average proliferation rate measured in each single animal, pooled with other animals in the same treatment arm to generate a mean per group. For the evaluation of proliferation index, high magnification images were taken with a 63x oil objective with the ApoTome 2 system (Carl Zeiss MicroImaging Gmbh, Germany), in two distinct regions of interest (ROI; 382 μm x 70 μm) at the border of tumor mass, one in the dorsal and one in the ventral part of the cerebral cortex. Cells labelled for Ki67 and/or Hoechst were counted using the Zen Blue Edition software (Carl Zeiss MicroImaging GmbH, Germany). The proliferation index was calculated as the proportion of Ki67-positive cells over the total sample of Hoechst-labelled cells in the selected ROIs for each coronal slice.

### Electrophysiology

In order to chronically record visual evoked potentials (VEPs), a bipolar electrode was implanted in the visual cortex (i.e. 3mm lateral to lambda, at the intersection between sagittal- and lambdoid-sutures) of C57BL/6J glioma-bearing mice with the tumor in the visual cortex as described(Cerri et al., 2016). A metal post was also placed on the occipital bone and fixed with dentistry cement (KERR, Dental Leader, Italy). After a period of habituation to the apparatus, awake head-fixed animals were subjected to recording sessions three times a week, from day 8 after tumor implantation until the visual response was detectable. VEPs were recorded in response to abrupt reversal (1Hz) of a horizontal square wave grating (spatial frequency 0.06 c/deg; contrast 30%), presented on a monitor (Sony; 40×30 cm; mean luminance 15 cd/m2). For each recording session, VEP amplitude was quantified by measuring the peak-to-trough amplitude as previously described(Pizzorusso et al., 2002; Porciatti et al., 1999; Restani et al., 2009; Vannini et al., 2016).

### Temozolomide administration and visual stimulation

Temozolomide (TMZ, Sigma Aldrich, USA) was dissolved in a solution of 25% of Dimethyl Sulfoxide (DMSO, Sigma Aldrich, USA) made in saline. Two groups of glioma-bearing mice implanted with chronic electrodes were daily weighed and injected with TMZ (40 mg/kg, i.p.) from day 12 after tumor induction until the last recording session. During the treatment, animals were regularly monitored according with the Mouse Grimace Scale, MGS (Langford et al., 2010) and sacrificed when they lost more than 20% of weight. The TMZ+VS group received a daily administration of TMZ plus visual stimulation, according to the stimulation protocol described above. For each grafting session, mice were randomly allocated to an experimental group (control, TMZ or TMZ+VS).

### Statistical data analysis

Data shown are mean ± s.e.m. Student’s *t*-test, or One-way ANOVA with appropriate post-hoc tests were used as detailed. For the comparison of the fraction of animals with detectable VEPs, Log-rank test was used.

Statistical analyses were conducted using GraphPad Prism 6 software (USA).

## RESULTS

### Neuronal processes infiltrate the glioma mass

We first examined the structural interactions between glioma cells and adjacent neurons. GL261 cells were transplanted into the visual cortex of C57BL/6J mice. At day 14, the tumor mass was clearly visible in the injected hemisphere (**Fig. 1A**). Coronal brain sections were stained for the neural marker NeuN and the axonal marker Tau (**Fig. 1B-D**). We found significant axonal staining within the glioma mass, indicating a physical interaction between glioma cells and neuronal processes (**Fig. 1B-D**).

**Figure 1.**
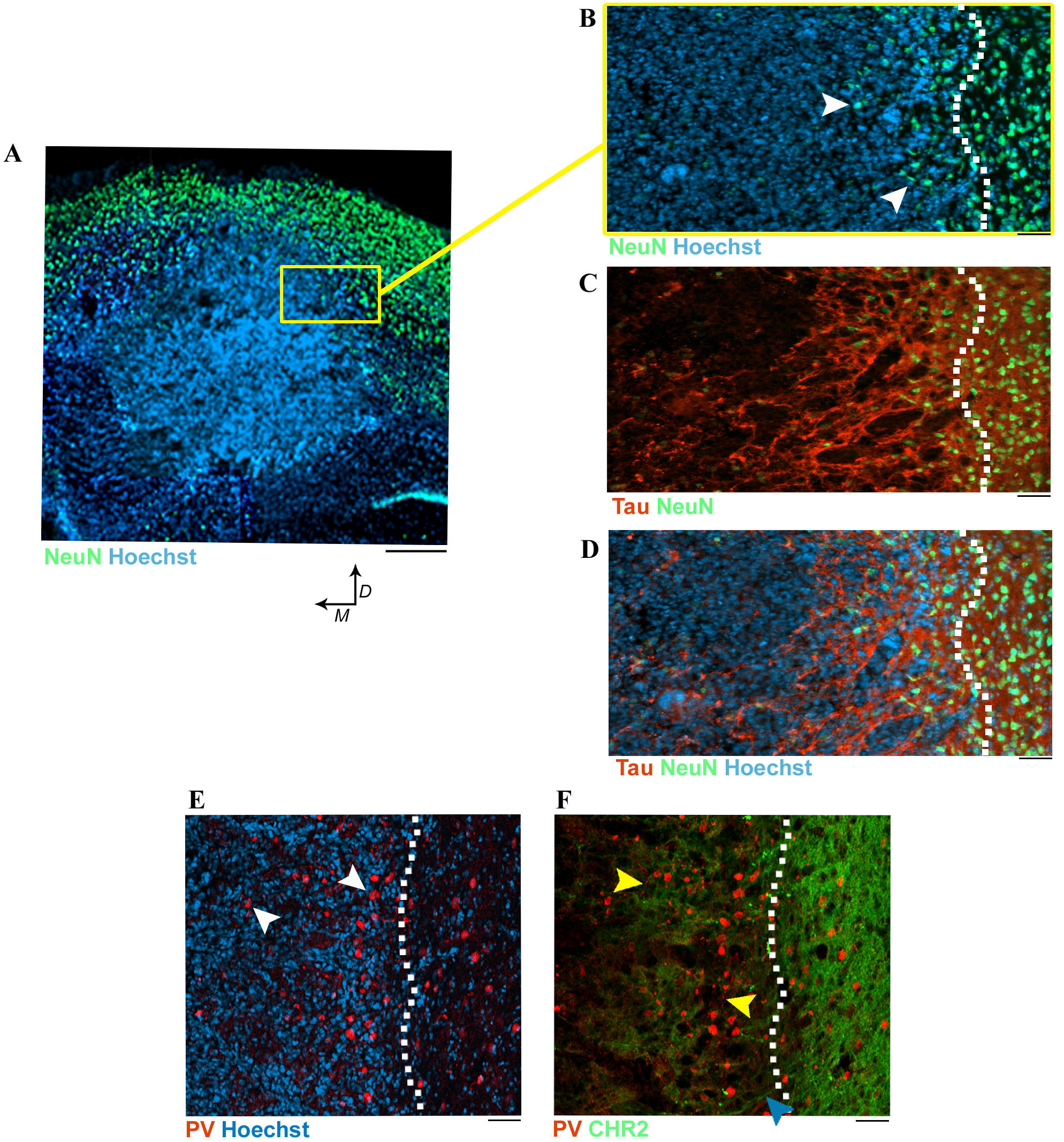
Neuronal processes penetrate the glioma mass. **A)** Representative tumor mass (Hoechst staining, blue) 14 days after GL261 cell transplant into the primary visual cortex. D, dorsal; M, medial. NeuN labelling is in green. Scale bar, 200 μm. **B-D)** High magnification images of a region at the border of the tumor mass. Dotted lines represent the presumptive tumor border. Scale bars, 100 μm. **B)** Neurons (in green; arrowheads) at the border of glioma mass (blue). **C)**Immunostaining for the axonal marker Tau (red) and NeuN (green) showing axonal fibers and neurons infiltrating the tumor. **D)** Triple labelling Tau (red)-NeuN (green) – Hoechst (blue). **E-F)** Neuroanatomical analysis of the tumor after inoculation of GL261 glioma cells in Thy1-ChR2-EYFP mice. Dotted lines represent tumor borders. Scale bars, 100 μm. **E)**Parvalbumin (PV) interneurons (red, arrowheads) inside the glioma mass (blue). **F)** Triple staining with Hoechst dye (blue), PV (red) and EYFP (green). Note processes of Parvalbumin interneurons (indicated by yellow arrowheads) and excitatory neurons (cyano arrowhead) within the border of glioma.

We next investigated whether both excitatory and inhibitory neurons contribute to neural infiltration within the tumor mass (**Fig. 1E-F**). We inoculated GL261 glioma cells into the visual cortex of Thy1-ChR2-EYFP mice, which express the light-gated cation channel channelrhodopsin-2 and enhanced yellow fluorescent protein selectively in excitatory, pyramidal neurons. Brain sections were also stained for parvalbumin (PV), a marker of fastspiking, GABAergic interneurons (Deidda et al., 2015).The neuroanatomical analysis detected processes and cell bodies of parvalbumin interneurons and excitatory neurons at the borders of the tumor (**Fig. 1E-F**).

### Optogenetic stimulation of pyramidal, excitatory neurons promotes glioma growth

Previous studies have shown that activation of cortical excitatory neurons through a 20 Hz optogenetic stimulation increases tumor proliferation in immunodeficient mice bearing patient-derived orthotopic glioma xenografts (Venkatesh et al., 2015, 2017). Here, we reproduced the single stimulation protocol after inoculation of GL261 cells in Thy1-ChR2-EYFP transgenic mice (**Fig. 2A, B**) (Spalletti et al., 2017).

**Figure 2.**
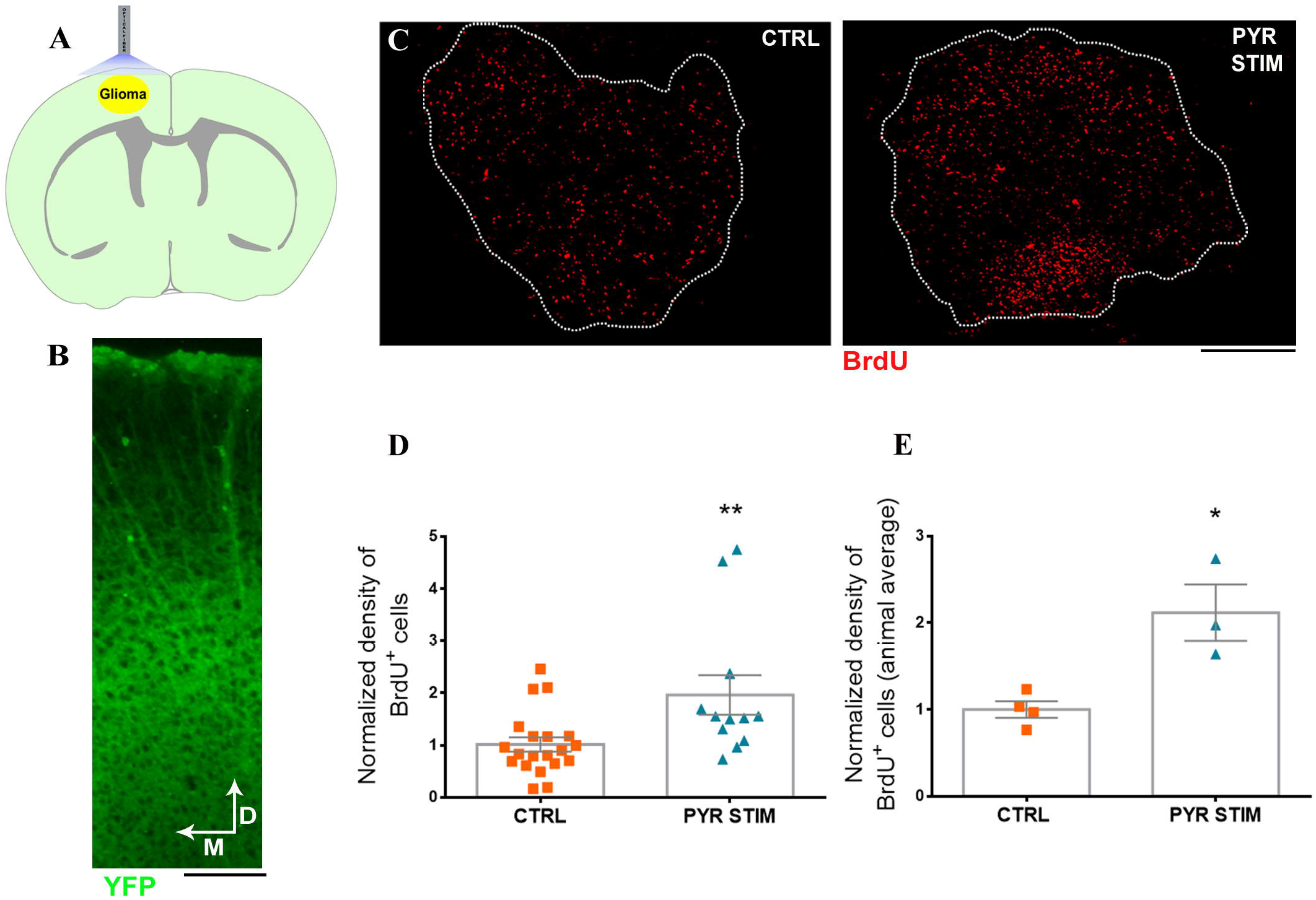
Optogenetic stimulation of pyramidal neurons promotes glioma proliferation. **A)** Schematic brain section showing the location of the tumor in motor cortex and the optic fiber for optogenetic stimulation in vivo. The green shading indicates widespread expression of ChR2 in pyramidal neurons of the transgenic Thy1-ChR2-EYFP mouse. **B)** Coronal section through the motor cortex showing expression of ChR2-EYFP (green) mainly in layer V. Scale bar, 200 μm. **C)** Representative staining for BrdU (red) in the motor cortex, 14 days after GL261 injection. CTRL, control with sham stimulation; PYR STIM, mouse with optogenetic stimulation of pyramidal neurons. The dotted lines indicate tumor borders. Scale bar, 200 μm. **D, E)** Normalized fraction of tumor area occupied by BrdU-positive cells in control (CTRL; n=20 slices from 4 animals) and stimulated mice (PYR STIM; n=12 slices from 3 animals): data from single slices (**D**) and averaged per animal (**E**). The mean ± s.e.m is also shown in grey for each group (Student’s t-test, **p=0.009 and *p=0.012, for D and E, respectively).

Animals were injected with GL261 cells in the motor cortex, and randomized to receive either real or sham (i.e. lights off) stimulation (**Fig. 2A**). Proliferating cells were assessed by BrdU incorporation (**Fig. 2C**). Quantitative analysis of the density of BrdU-labelled cells showed that the 20 Hz optogenetic stimulation of pyramidal neurons significantly increased tumor proliferation (Student’s *t*-test, p < 0.009 and p < 0.05; **Fig. 2D** and **2E**), confirming the results described in literature(Venkatesh et al., 2015).

### Optogenetic stimulation of Parvalbumin-positive, GABAergic interneurons restrains glioma proliferation

We next asked whether the activity of PV-positive, fast-spiking interneurons has an impact on tumor proliferation. PV-Cre mice (Cardin et al., 2009; Mariotti et al., 2018) were inoculated into the motor cortex with a AAV vector carrying the doublefloxed ChR2-mCherry sequence. The neuroanatomical analysis indicated that over 80% of the PV cells were positive for mCherry in the transduced area (**Fig. 3A, B**). One week later, mice were injected with GL261 cells in the motor cortex, and they were randomized into two groups, to receive either real or sham (i.e. lights off) stimulation. A quantitative analysis of the density of proliferating cells was performed by both Ki67 (**Fig. 3C**) and BrdU immunolabeling. The data clearly showed that 40 Hz optogenetic stimulation of PV interneurons significantly restrains tumor proliferation. Quantification of Ki67- and BrdU-positive cells indicated dampened proliferation, considering either single slices (Student’s *t*-test, p < 0.001; **Fig. 3D, E**) or the average value for each animal (Student’s *t*-test, p < 0.05; **Fig. 3F, G**). Thus, selective stimulation of fast-spiking interneurons reduces the extent of proliferating cells in the tumour mass.

**Fig 3.**
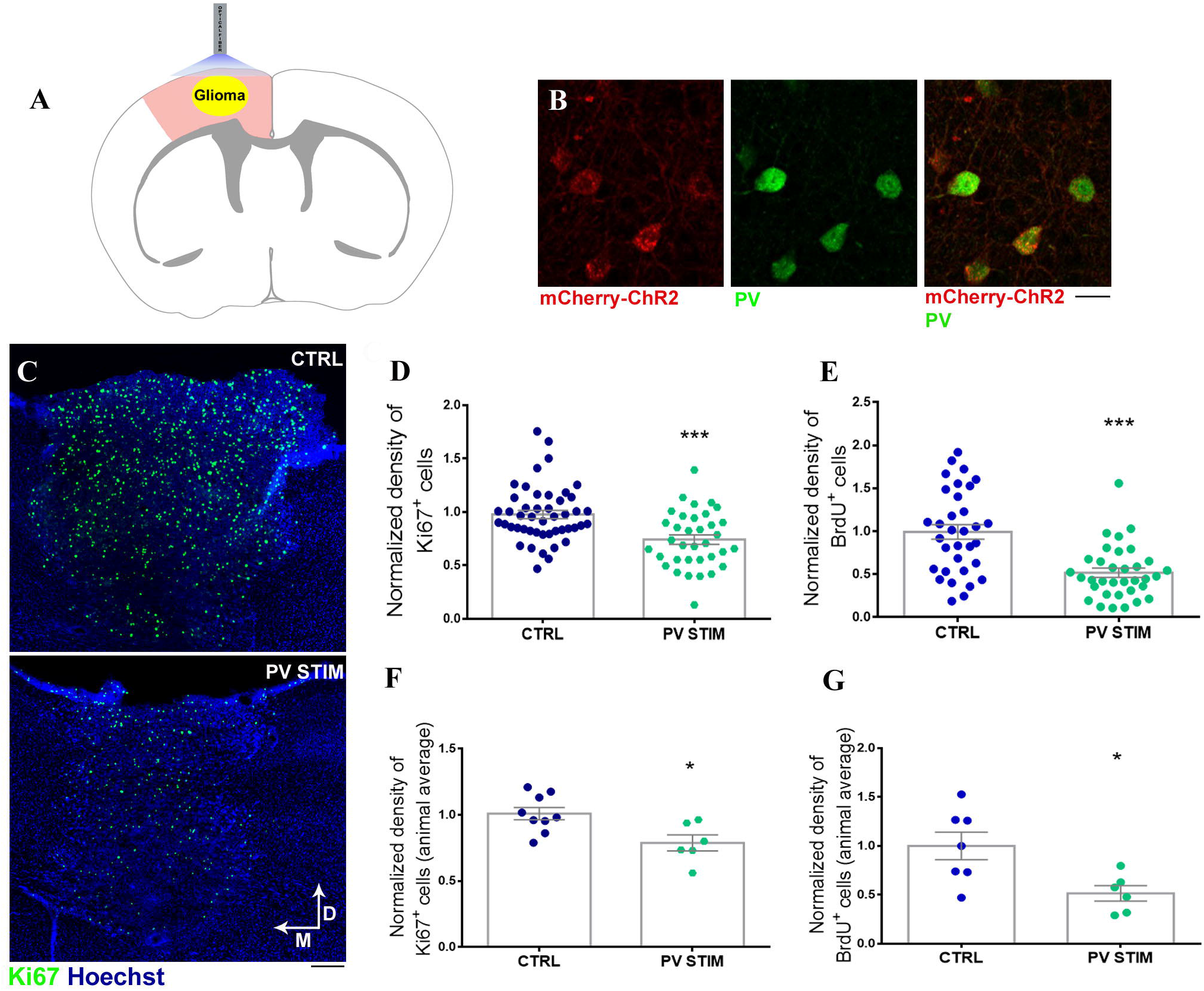
Optogenetic stimulation of Parvalbumin interneurons inhibits cell proliferation in tumour mass. **A**) Schematic cartoon illustrating the optogenetic stimulation of PV interneurons in a coronal section of the motor cortex. The red shading in the cortex indicates localized transduction of PV interneurons with the AAV vector. **B**) Representative images of a cortical section in a PV-Cre mouse transduced by AAV carrying mCherry-ChR2. The section is double labelled for mCherry (red) and Parvalbumin (green). Colocalization is shown in the panel at right. Scale bar, 50 μm. **C**) Representative images of proliferating Ki67-positive cells (green) within the glioma mass (blue) in the motor cortex of sham (CTRL, top) and stimulated (PV STIM, bottom) mouse 14 days after glioma injection. M, medial; D, dorsal. Scale bar, 100 μm. **D-G**) Normalized fraction of tumor area occupied by Ki67- (**D, F**) and BrdU- (**E, G**) positive cells in control (CTRL: Ki67, n=47 slices from 9 animals; BrdU, n=33 slices from 6 animals) and stimulated mice (PV STIM: Ki67, n=35 slices from 6 animals; BrdU, n=35 slices from 6 animals). Data from single slices (**D, E**) and averaged per animal (**F, G**). The mean ± s.e.m is also shown in grey for each group (Student’s t-test, ***p<0.001, *p=0.012 for F, *p=0.015 for G).

### Afferent sensory input bidirectionally regulates tumor proliferation

Having established the distinct roles of pyramidal and GABAergic cells in the control of glioma growth, we next investigated the impact of a coordinated activation of excitatory and inhibitory neurons triggered by sensory experience. For this experiment, tumors were placed in the occipital cortex, as afferent sensory input can be easily downregulated via dark rearing (Gianfranceschi et al., 2003) or upregulated with controlled visual stimulation(Restani et al., 2009).

Accordingly, we implanted mice with GL261 cells in the primary visual cortex and all animals were maintained in a normal light/dark cycle until day 11. They were then randomized to remain in standard laboratory conditions (standard laboratory, SL), placed in total darkness (dark rearing, DR) or visually stimulated during the light period (visual stimulation, VS) by placing them for 8 hr daily in front of gratings of various contrasts as well as temporal and spatial frequencies (**Fig. 4A**). Stainings for cell proliferation were conducted on day 14.

**Figure 4.**
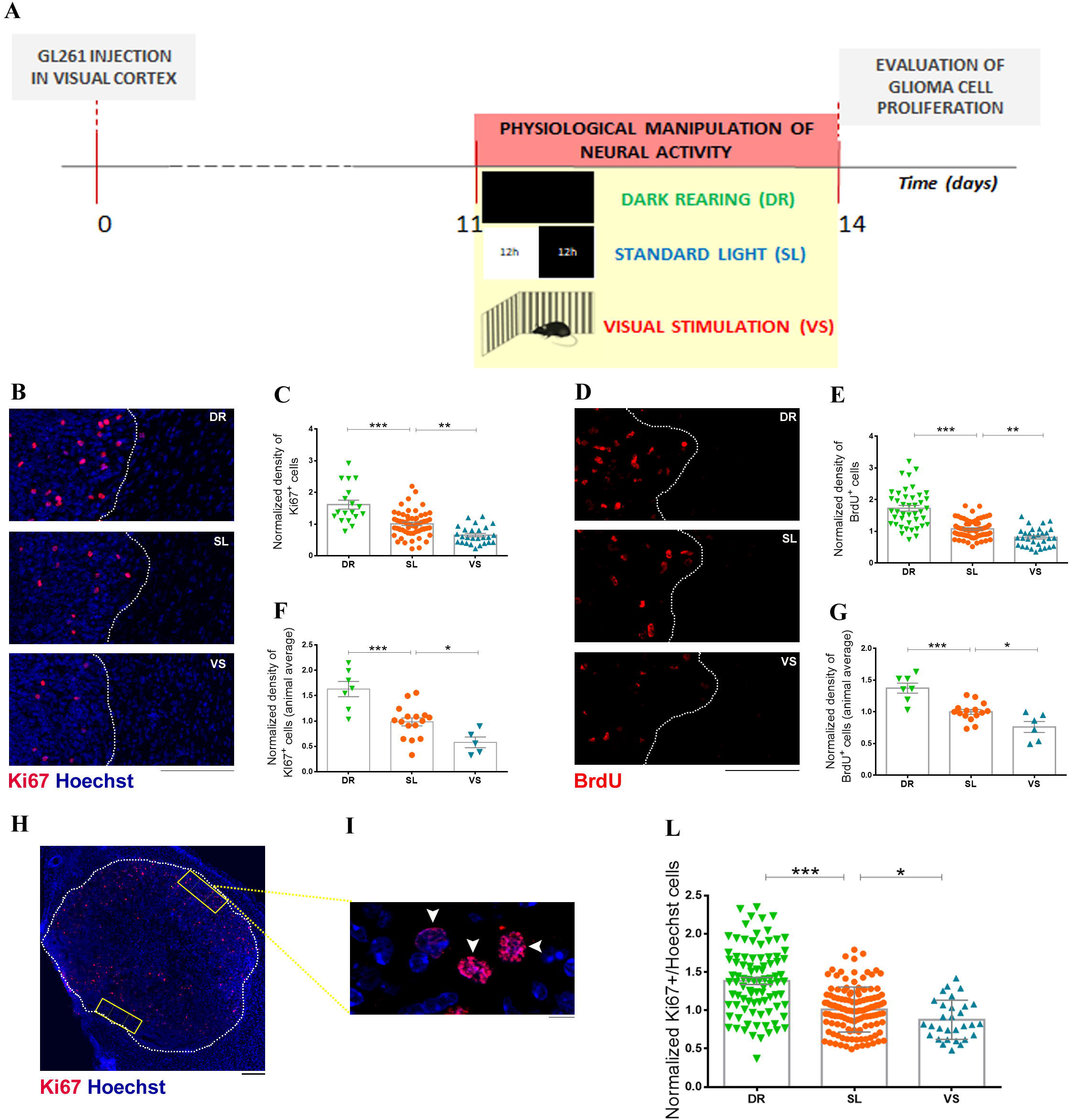
Manipulation of sensory input bidirectionally impacts glioma cell proliferation. **A)**Experimental protocol. **B, D**) Representative immunostainings for Ki67 (**B**) and BrdU (**D**) (red) within the tumor mass in the visual cortex of glioma-bearing mice, 14 days after GL261 cell transplant. Dotted lines represent tumor borders. DR, dark rearing; SL, standard light conditions; VS, visual stimulation. Scale bar, 200 μm. **C, E, F, G)**Normalized fraction of tumor area occupied by proliferating cells in the different experimental groups: data from single slices (**C,E**) and averaged per animal (**F,G**) (DR: Ki67, n=18 slices from 7 animals; BrdU, n=42 slices from 7 animals; SL: Ki67, n=58 slices from 16 animals; BrdU, n=63 slices from 15 animals; VS: Ki67, n=26 slices from 5 animals; BrdU, n=31 slices from 6 animals). The mean ± s.e.m is shown in grey for each group. (*p < 0.05, **p < 0.0025, ***p < 0.001). **H**) Representative immunostaining for Ki67 showing proliferating cells (red) at the border of tumor mass in the visual cortex. Blue: Hoechst counterstaining. Two ROI (regions of interest) for the cell count at the border of the tumor mass are indicated in yellow. Scale bar, 200 μm. **I**) High magnification of dividing (arrowhead) and non-dividing (blue) cells in a ROI. Scale bar, 10 μm. **L**) Proliferation index indicating the fraction of Ki67-positive cells with respect to all Hoechst-stained nuclei (DR, n=90 slices from 6 mice; SL, n=125 slices from 7 mice; VS, n=30 slices from 3 mice). The mean ± s.e.m is shown in grey for each group. (*p < 0.05, ***p < 0.001).

We found that dark rearing enhanced while visual stimulation dampened the density of proliferating cells in the tumor mass (**Fig. 4B** and **4D**), indicating a bidirectional regulation of glioma proliferation by afferent sensory input. As in the previous experiments, we first performed the statistical analysis by cumulating the data obtained from all the brain sections within each experimental group (for Ki67 and BrdU: One-way ANOVA followed by Dunnett’s test, DR vs SL p<0.001, VS vs SL p<0.0025; **Fig. 4C** and **4E**). Second, we performed the analysis on a per mouse basis, i.e. using the average densities measured in the animals reared in a specific condition (for Ki67 and BrdU: One-way ANOVA followed by Dunnett’s test, DR vs SL p<0.001, VS vs SL p<0.05; **Fig. 4F** and **4G**).

To further corroborate these results, we calculated the proliferation index at the border of the tumor mass on all the analyzed coronal slices, by counting the proportion of tumor cells expressing Ki67 over the total number of Hoechst-stained nuclei in each region of interest (**Fig. 4H, I**). Also in this case, whereas light deprivation by DR increased the number of dividing cells, visual stimulation reduced the proliferation index with respect to control glioma-bearing animals reared in standard light conditions (SL) (Student’s t-test, DR vs SL p<0.001, VS vs SL p<0.05; **Fig. 4L**). Altogether, these data demonstrate that levels of afferent sensory input bidirectionally regulate tumor growth in the GL261 glioma model.

### Effects of sensory input on tumor proliferation are region-specific

The data reported above demonstrate that sensory stimulation modulates glioma cell proliferation. To determine whether these effects remain localized to the visual cortex, or involve secreted factors which may act at distance, animals were implanted with GL261 glioma cells into the motor cortex and then subjected to visual deprivation via dark rearing (**Fig. 5A**) starting from day 11 after tumor injection. A second group of animals with glioma in the motor cortex were reared in standard light conditions. The quantitative analysis showed no significant differences between deprived (DR) and standard light (SL) mice in the proliferation rates, measured via the density of both Ki67- and BrdU-positive cells (Student’s t-test, p>0.05; **Fig 5B-E**). We conclude that the effect of sensory input on glioma cell proliferation is region-specific.

**Figure 5.**
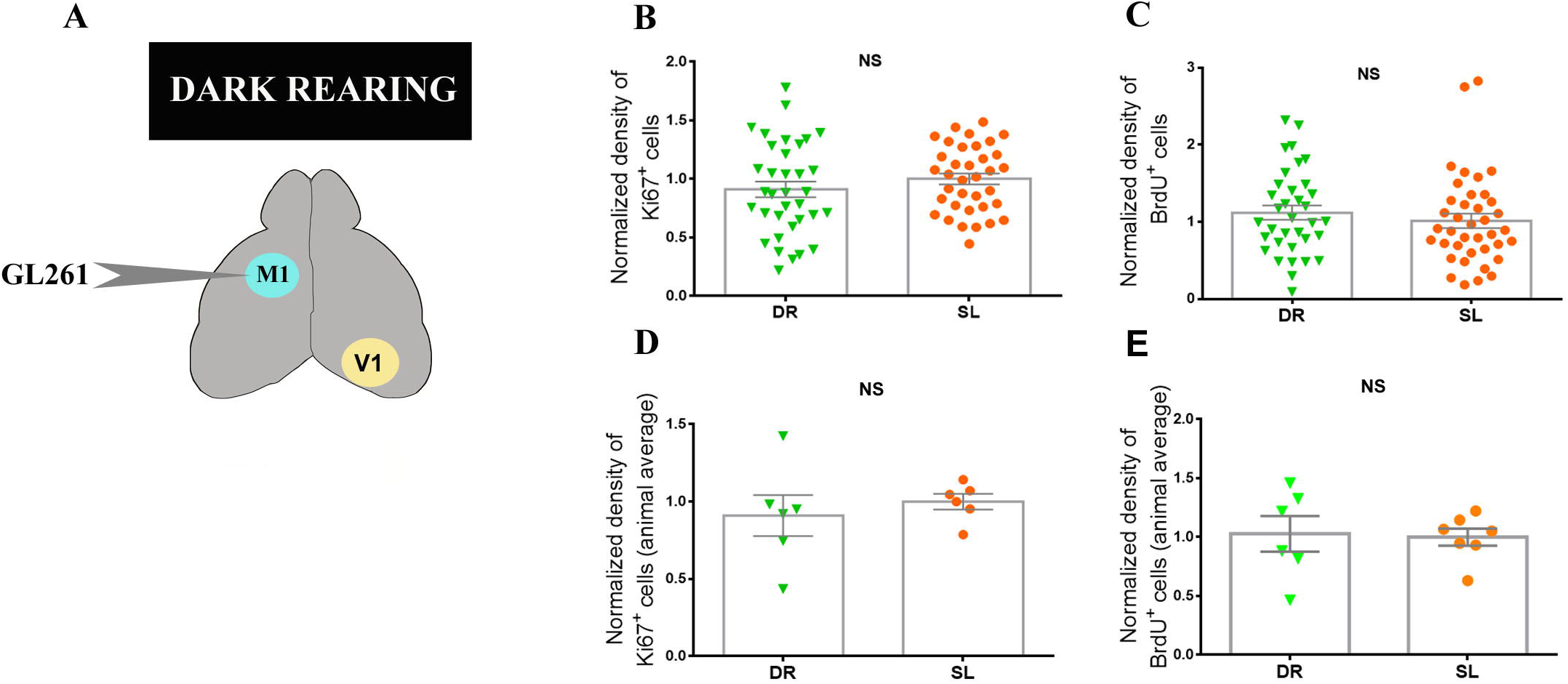
Dark rearing has no impact on tumor proliferation in motor cortex. **A)**Schematic drawing of the experimental protocol: mice injected with GL261 cells in the primary motor cortex (M1) were reared in total darkness. V1, primary visual cortex. **B, C**) Normalized fraction of tumor area occupied by proliferating Ki67^+^ (**B**) and BrdU^+^ (**C**) cells (Ki67: DR, n=35 slices from 7 animals; SL, n=36 slices from 6 animals; BrdU: DR, n=35 slices from 6 animals; SL, n=39 slices from 7 animals). **E, F)** Average values for each animal are reported. The mean ± s.e.m is shown in grey for each group (p > 0.05). NS, not significant.

### Sensory stimulation combined with temozolomide treatment delays the deterioration of visual responses induced by glioma growth

We next tested whether sensory stimulation can be exploited as an adjuvant to classical chemotherapy (i.e., temozolomide – TMZ) to delay neurological dysfunction in peritumoral neurons. As a readout of neuronal responses, we used longitudinal measurements of visual evoked potentials (VEPs) in mice bearing tumors in the occipital cortex.

Animals were injected with GL261 cells into the visual cortex and implanted with a bipolar electrode to record VEPs from day 8 post injection (baseline recording). Typical VEP responses consisted or an early negative wave and a late positive peak with latencies of approx. 60 and 110 ms, respectively (**Fig. 6A**). Glioma-bearing mice were either left untreated (n = 6) or administered TMZ daily starting from day 12 after tumor inoculation (TMZ, n = 7). Another cohort of TMZ animals received also daily visual stimulation for 8 hours (TMZ+VS, n = 9). After each grafting session, inoculated animals were randomly allocated to the three groups.

**Figure 6.**
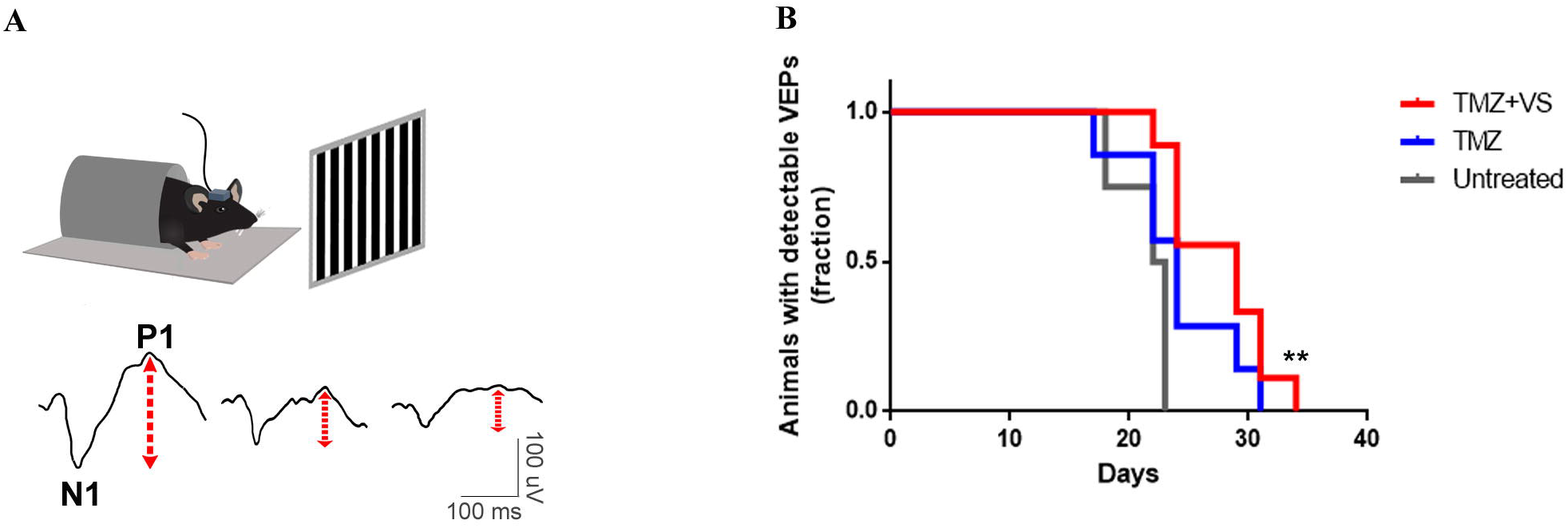
Visual stimulation combined with TMZ administration delays the impairment of cortical responses due to glioma progression. **A)** (Top) Schematic drawing of the experimental set-up for chronic VEP recordings in awake, head-restrained mice bearing GL261 cells in the visual cortex. (Bottom) Representative examples of VEP recordings at day 8 (left), 12 (middle) and 19 (right) following tumor inoculation. Note clear decrease of VEP amplitudes during tumor progression. N1, early negative peak; P1, positive peak. **B)** Fraction of animals with detectable VEPs in glioma-bearing animals that were left untreated (grey line, n = 6), administered with TMZ (blue line, n = 7), or given TMZ plus visual stimulation (TMZ+VS, red line, n = 9; **p=0.0023 with respect to untreated group).

A progressive deterioration of the visual response was observed in all glioma-bearing mice (**Fig. 6A**), but the groups showed different kinetics. Temozolomide monotherapy slightly protracted the time window of visual responsiveness, but the effect was not statistically significant (Log-rank test, p=0.17). The evaluation of VEP amplitudes demonstrated the maintenance of a detectable visual responsivity of peritumoral neurons in TMZ+VS group (Log-rank test, p=0.0023; **Fig. 6B**). Thus, a therapy combining TMZ and visual stimulation delays the loss of neural responses in glioma-bearing mice.

## DISCUSSION

Recent experiments have provided strong evidence for a bidirectional communication between glioma cells and adjacent neurons. Release of glutamate from glioma cells induces seizures and excitotoxicity in peritumoral networks; glutamate also acts in a paracrine/autocrine manner on glioma cells, facilitating their spread into the brain parenchyma (de Groot and Sontheimer, 2011). Genes encoding glutamate receptors are expressed in a subset of glioma cells, and excitatory functional synapses are established between neurons and cancer cells (Venkataramani et al., 2019; Venkatesh et al., 2019). Optogenetic stimulation of pyramidal neurons in the tumor-adjacent zone triggers calcium signals in glioma cells, increasing their proliferation and invasiveness (Venkataramani et al., 2019; Venkatesh et al., 2019).

In addition to principal cells, peritumoral cortical circuitry also comprises GABAergic interneurons. The largest class (40-50%) of GABAergic interneurons is represented by fastspiking, parvalbumin-positive cells. They are distributed throughout all cortical layers, and form synapses on the soma and proximal dendrites of pyramidal cells and other interneurons (Lim et al., 2018). In the peritumoral zone, an overall loss of peritumoral fast spiking interneurons and a reduction of their firing rates has been described (Tewari et al., 2018). This reduced firing is due to the degradation of perineuronal nets that surround these interneurons, and is critically involved in tumor-associated epileptic seizures (Tewari et al., 2018). So far, no data were available on the role for these GABAergic cells in tumor progression.

In this manuscript, we used two specific transgenic lines to perform selective optogenetic stimulation of pyramidal and fast-spiking interneurons in the tumor-adjacent zone of glioma-bearing mice. This has allowed us to compare directly the impact of the activity of principal cells and interneurons on tumor proliferation.

We employed intracortical transplants of GL261 cells which are widely used in experimental glioma research. One advantage is that the GL261 system represent one of the best characterized syngeneic, immunocompetent models(Oh et al., 2014). Limitations include the fact that this model is not molecularly faithful to human GBM (it is driven by a set of mutations not observed in human GBM) and typically grows as a large solid mass rather than a diffusely invasive cancer. Despite this growth pattern, we have demonstrated robust infiltration of neuronal processes within the tumor mass (**Fig. 1**), which represent the structural basis for interactions between tumor cells and the neuronal microenvironment. Recent studies have shown that nodular tumors, such as those observed in GL261 model, cause compression and dysfunction of nearby brain tissue (Seano et al., 2019). A direct molecular communication between peritumoral neurons and GL261 cells has been described(Liu et al., 2013). The authors suggest that peritumoral neurons restrain glioma progression through the pivotal role of PD-L1 protein, and indeed neuronal PD-L1 signaling in brain cells is important for GBM patient survival (Liu et al., 2013).

### Optogenetic stimulation of principal cells and fast-spiking interneurons has opposite effects on tumor proliferation

The optogenetic experiments were clear in indicating that pyramidal cell stimulation enhances cell proliferation in tumour mass, consistent with several previous reports (Venkatesh et al., 2015, 2017). The selective stimulation of parvalbumin-positive interneurons elicited the opposite effect – i.e. a significant reduction of glioma cell proliferation. Thus, glutamatergic and GABAergic signalling appear to have opposite actions on tumor growth. Similar antagonistic effects of neurotransmitter systems have been described in breast cancer, where stimulation of parasympathetic (cholinergic) afferents reduces tumor growth, while stimulation of sympathetic (noradrenergic) nerves accelerates cancer progression (Kamiya et al., 2019). The effects of stimulation of parvalbumin interneurons is consistent with the finding that human glioblastoma cells express functional GABA-A receptors and that endogenous GABA released by tumor cells attenuates proliferation (Blanchart et al., 2017). In addition to the autocrine/paracrine effects of tumor-derived GABA, the present results indicate that peritumoral interneurons may represent a further source of GABA, supporting anti-proliferative effects. The high vulnerability of fast-spiking interneurons to glioma-induced excitotoxicity in the tumor-adjacent zone (Tewari et al., 2018) may trigger a vicious cycle with reduced GABA release, increased firing of pyramidal neurons, and consequently enhanced tumor growth. A similar scenario is indicated by the recent work of Deneen and collaborators (Yu et al., 2020), where specific mutations harboured by transplanted glioma cells render tumours more aggressive and lethal. Gliomas driven by these variants shape synaptic connectivity in the peritumoral region, with enhanced density of excitatory boutons and decreased inhibitory synapses, and consequently heightened network hyperexcitability. These data concur in indicating that glioma-derived factors may drive synaptic changes which result in aberrant activity patterns leading to increased tumor growth.

### Afferent sensory input modulates tumor proliferation

Pyramidal neurons and fast-spiking interneurons are directly activated by sensory input via thalamocortical afferents (Sugiyama et al., 2008; Yu et al., 2016). Therefore we wondered whether and how sensory stimulation affects glioma cell proliferation. To address this issue, we exploited glioma cell grafting in the visual cortex, a region where afferent activity can be easily modulated (Deidda et al., 2015; Porciatti et al., 1999; Vannini et al., 2016). While high-grade gliomas in the visual cortex are rare, about 30-50% of the cortical mantle in humans is responsive to visual stimulation, making the results of this experiential modulation of sensory input potentially interesting in a translational perspective. Specifically, we employed either visual stimulation or dark rearing for 3 consecutive days to transiently enhance or reduce sensory input to the peritumoral areas. The proliferation analyses indicated that dark rearing increases glioma cell proliferation, while stimulation with gratings of different spatial and temporal frequencies has the opposite effect, counteracting tumor cell division. We chose to employ short periods of visual stimulation/deprivation as activity in occipital areas is known to homeostatically rescale within one week to altered levels of visual input (Turrigiano, 2012). For example, in the case of chronic visual deprivation, pyramidal cell firing in the visual cortex is initially depressed during the first 2-3 days but then returns to baseline at one week due to homeostatic compensation (Hengen et al., 2016; Mrsic-Flogel et al., 2007).

Activation of the cortical network by sensory stimuli results in a balanced and sequential activation of several classes of pyramidal cells and interneurons, in a layer-specific manner(Kerlin et al., 2010; Okun et al., 2015). Can the impact of visual stimulation be explained based on a differential involvement of principal and GABAergic neurons? There are several possibilities to be considered. First, while pyramidal neurons show exquisite selectivity for properties of the visual stimulus (spatial and temporal frequency, contrast), fast-spiking interneurons (and other GABAergic cells) are broadly tuned for stimulus features (Kerlin et al., 2010). This means that during exposure to a specific grating, only a subset of pyramidal neurons are firing while GABAergic interneurons display a more global activation. Second, in the awake mouse, cortical responses to visual stimulation are precisely shunted by GABAergic inhibition, restricting the spatial spread and temporal persistence of pyramidal cell discharges (Haider et al., 2013). Third, parvalbumin-positive interneurons fire at much higher rates than cortical pyramids. Altogether, these data suggest that visual stimulation may trigger substantial GABA release that overcomes the effect of glutamatergic signalling.

After dark rearing, a substantial decrease of GABA-immunoreactive neurons has been found in primary visual cortex (Benevento et al., 1995). This reduction of GABAergic cells may remove a neurochemical brake to tumor growth, thus explaining the higher proliferation rates following visual deprivation. Importantly, the impact of visual input is region specific, as shown by the lack of effect of dark rearing in mice with syngeneic glioma in the motor cortex.

One important goal of glioma therapies is the preservation of function in the tumor-adjacent zone (Vannini et al., 2016, 2017). For example, subjects with tumors in the occipital regions of the brain show several types of visual impairments that potently impact their quality of life (Sharrack et al., 2018). To study the decay of visual function in glioma-bearing mice, we set up a technique of chronic VEP recordings to longitudinally follow the functionality of visual cortex during tumor progression. We combined TMZ delivery with daily visual stimulation, as an addon of established therapy for GBM. In the TMZ+visual stimulation group, we found a significant prolongation of neural responsiveness, suggesting a possible strategy to delay the loss of sensory functions during glioma growth.

In summary, the data reported in this manuscript demonstrate a key and previously unrecognized role for parvalbumin-positive, GABAergic neurons in controlling glioma cell proliferation. We also show that levels of sensory afferent input bidirectionally modulate tumor proliferation. Agents that stimulate GABAergic signalling may be tested to improve the efficacy of conventional anti-glioma treatments(Laub et al., 2018; Wick et al., 2018).

## Author contributions

Experimental design, implementation of experiments, analysis and interpretation of the data: E.T., E.V., C.S., A.C.

Experimental design, analysis and interpretation of the data: C.C., M.Co., M.Ca. Interpretation of the data: C.M.M.

Writing and critical revision of the manuscript: E.T., E.V., C.C., M.Co., M.Ca.

## Acknowledgements

We thank Francesca Biondi (CNR Pisa) for excellent animal care. E.V. and C.C. were supported by postdoctoral fellowships from Fondazione Umberto Veronesi, Milan, Italy.

